# NF kappa B Regulator Bcl3 Controls Development and Function of Classical Dendritic Cells in *Toxoplasma gondii* Infection

**DOI:** 10.1101/2022.04.07.487586

**Authors:** June Guha, Byunghyun Kang, Estefania Claudio, Neelam R. Redekar, Hongshan Wang, Brian L. Kelsall, Ulrich Siebenlist, Philip M. Murphy

## Abstract

The atypical IκB family member Bcl3 associates with p50/NF-κB1 or p52/NF-κB2 homodimers in the nucleus, and positively or negatively modulates transcription in a context-dependent manner. In mice lacking Bcl3 globally or specifically in CD11c^+^ cells, *Toxoplasma gondii* infection is uniformly fatal and is associated with an impaired Th1 immune response. Since Bcl3 expression in dendritic cells (DC) is pivotal for antigen presentation and since classical DCs (cDC) are major antigen presenting cells, we investigated the role of Bcl3 specifically in cDCs in *T. gondii* infection *in vivo* by crossing Zbtb46 cre mice with *Bcl3^flx/flx^* mice. The conditional cDC Bcl3 KO was as susceptible to lethal *T. gondii* infection as the total Bcl3 KO and generated poor Th1 responses. Splenocyte single cell RNA seq in the model revealed defective Bcl3-dependent expression of genes involved in antigen processing. Consistent with this, soluble toxoplasma antigen presentation was impaired in Bcl3-deficient cDCs, and tetramer staining demonstrated defective *T. gondii* antigen-specific splenic CD4^+^ and CD8^+^ T cell responses in infected cDC *Bcl3^-/-^* mice. *In vitro* differentiation of bone marrow progenitors from wildtype and cDC *Bcl3^-/-^* mice using Flt3L, NOTCH and IFN-γ stimulation recapitulated the defective Bcl3-dependent cDC antigen-presentation activity observed in vivo. Splenocyte single cell RNA seq also revealed the existence of a unique subpopulation of Zbtb46^+^LysM^+^ DC which exhibited Bcl3-dependent expansion after infection. We also detected cDCs coexpressing the monocytic markers CD64 and Ly6C (designated icDC1 and icDC2) mainly in infected spleen, which were less abundant in *Bcl3^flx/flx^ Zbtb46 cre* mice. Together, our results indicate that Bcl3 in classical DCs is a major determinant of protective T cell responses and survival in *T. gondii*-infected mice, and shapes DC ontogeny.

**Author Summary:** Dendritic cells initiate immune responses against invading pathogens. As professional antigen presenting cells they process and present antigen via the major histocompatibility complex to T cells and thus activate them. Bcl3, an atypical member of the IκB family regulates the APC function of dendritic cells. In this study we show that expression of Bcl3 specifically in classical DCs is critical for host protection against a protozoan parasite, *Toxoplasma gondii*. Host protective proinflammatory mechanisms are compromised in mice deficient in Bcl3 in classical DCs leading to an elevated organ parasite load and eventually death of the infected animals. We also found the emergence of Bcl3-dependent hybrid DCs upon *T. gondii* infection, which have mixed phenotypic markers from DCs and monocytes. Antigen processing genes are significantly downregulated in Bcl3-deficient cDCs, which may account for defective cross presentation of *T. gondii* antigens. In an in vitro differentiation model, we showed that development of XCR1^+^cross presenting cDC1s is critically regulated by Bcl3. Overall, this study reveals the complexity of dendritic cell ontogeny and the role of Bcl3 in classical DC function in the context of *Toxoplasma* infection.

## Introduction

The NF-κB family of transcription factors acts as a master regulator of diverse physiological processes, from cell survival and proliferation to inflammatory responses against environmental stimuli and infectious agents. It consists of 2 subfamilies, Rel/NF-κB and IκB (inhibitor of κB). The Rel/NF-κB subfamily members include Rel A, Rel B, c-Rel, p50 and p52, which form homo- or heterodimers and are able to modulate transcription of target genes by binding to κB enhancer elements [1]. The IκB subfamily regulates NF-κB function and is divided into two subgroups, the classical (IκBα, IκBβ, IκBε, p100 and p105) and atypical IκB proteins (Bcl3, IκBζ and IκB NS). Classical IκB proteins inhibit NF-κB dimers through direct interactions in the cytoplasm. Following cell activation, classical IκB proteins become phosphorylated and subjected to ubiquitin-mediated degradation which releases NF-κB dimers for translocation to the nucleus to activate target genes [2]. In contrast, atypical IκB proteins do not undergo activation-dependent cytoplasmic degradation and instead modulate transcription in the nucleus [3].

Bcl3 (B cell lymphoma factor 3) was originally identified as a gene involved in genomic translocations in cases of B cell chronic lymphocytic leukemia (B-CLL) [4]. Bcl3 preferentially transactivates p50 or p52 homodimers by interactions involving its ankyrin domains; however, it may inhibit or stimulate transcription of NF-κB target genes in a highly context-dependent manner [5, 6]. Analysis of Bcl3 knockout mice has revealed diverse immunoregulatory roles, for example, in T and B lymphocyte development [7, 8], proper formation of splenic architecture [9], terminal differentiation of memory CD8^+^ T cells [10] and dendritic cell function [11]. Accordingly, Bcl3 deficient mice have been reported to have increased susceptibility to infectious agents, including *Klebsiella pneumoniae, L. monocytogenes, S. pneumoniae* and *Toxoplasma gondii.* [12, 13].

*Toxoplasma gondii* is an opportunistic obligate intracellular protozoan and a member of the phylum Apicomplexa. It is capable of infecting almost all nucleated cells and can establish a long-term latent infection in the host. Toxoplasma has a complex life cycle in mammals, including a sexual reproductive cycle in definitive feline hosts, and an asexual cycle in intermediate hosts, which include humans. The clinical presentation is variable and depends on the immune status of the infected host. Immunocompetent individuals may develop a mononucleosis syndrome or remain asymptomatic, but in both cases go on to develop life-long latent infection. However, in immunocompromised hosts the parasite may reactivate resulting in toxoplasma encephalitis or retinochoroiditis. The infection may be particularly life-threatening to the fetus during pregnancy, resulting in congenital developmental abnormalities, including hydrocephalus, microcephaly, cerebral calcifications, retinochoroiditis, blindness, epilepsy, motor retardation, and anemia[14, 15] .

After infection, *T. gondii* replicates rapidly as a tachyzoite form by endogeny, then lyses the infected cell and spreads to neighboring cells. The organism can cross the blood brain barrier and ultimately become encysted in brain and skeletal muscle as a slowly replicating bradyzoite form resulting in latent infection [16]. *T. gondii* infection leads to a strong cell-mediated immune response. After transmigration across polarized epithelial cells, the parasite encounters dendritic cells which sense the pathogen and produce large amounts of IL-12, which activates NK cells to produce IFN-γ during the acute phase. Subsequently, parasite-specific Type 1 CD4^+^ T cells and cytotoxic CD8^+^ T cells produce IFN-γ as the infection enters a chronic phase [17]. Dendritic cells coordinate the immune response through antigen presentation and pro-inflammatory cytokine production, particularly IL-12, resulting in priming of CD4^+^ and CD8^+^ T cells for IFN-γ production.

Dendritic cell (DC) differentiation is highly organ-specific and depends on the inflammation status of the host. Hematopoietic progenitor cells differentiate into common myeloid and lymphoid progenitors (CMP and CLP) that give rise to precursors to classical DCs (cDCs) and plasmacytoid DCs (pDCs), respectively in the bone marrow. These precursors are released into the blood and subsequently seed both lymphoid and non-lymphoid tissues where they develop into mature cDCs and pDCs. cDCs are further divided into cDC1 and cDC2 populations based on expression of surface molecules and dependence on unique transcription factors (TFs) for their development. cDC1 are characterized largely as CD11c^+^MHC II^+^CD11b^-^ XCR1^+^ cells and express CD103 in peripheral tissues, and CD8αα in lymphoid tissues. cDC1s are dependent on the TFs Irf8, Batf3, Nfil3, and Id2. In contrast, cDC2s are largely CD11c^+^MHC II^+^CD11b^+^Sirpa^+^ and rely on Irf4 for their full development, but are more heterogeneous in that some express CD103 in the intestine, and subsets have been defined that rely on either Notch 2 or Klf4 for their differentiation [18, 19].

In nonlymphoid organs, 1-5% of cells are cDCs comprised of CD103^+^CD11b^-^ cDC1 and CD11b^+^ cDC2 subsets, although a minor population of CD11b^-^CD103^-^ cells is present in the intestine which remains less well defined. In the spleen and LNs, CD8^+^ cDC1s constitute 20-40% of total cDC, the rest being CD11b^+^ cDC2s and pDCs. cDC1 and cDC2 migrating in lymph from peripheral tissues (designated cDC1mig and cDC2mig) express CCR7 and are usually abundant in T cell zones of draining lymph nodes (LNs). They can usually be distinguished from resident DCs in the steady state by relatively higher MHCII and lower CD11c expression, however under inflammatory conditions, when resident DCs are activated, they can no longer be distinguished by these markers. Resident LN DCs on the other hand are derived from blood precursors and remain in organized lymphoid tissues [20] .

cDCs play critical roles in both innate and adaptive immunity. cDC1s regulate CD8^+^ and Th1 T cell responses against viruses and intracellular pathogens by providing high amounts of IL-12, and by cross-presenting antigen to naïve CD8^+^ T cells. cDC2s are critical for initiating immune responses against extracellular bacteria and fungi. They are thought to be highly capable of priming CD4^+^ T cells, produce IL-23, IL-6 and TGF-β which contributes to the polarization of Th17 cells [19], and in some contexts strongly drive the differentiation of Th2 cells. However, these functional attributes of cDC1 and cDC2 are somewhat plastic, particularly in infections and other inflammatory conditions, where cDC2s for example can cross-present antigen to CD8 T cells, and cDC1 are fully capable of presenting antigens to naïve CD4^+^ T cells. Finally, monocytes are also known to differentiate into inflammatory cells during infection and other inflammation which are capable of presenting antigens to CD4^+^ and CD8^+^T cells and have thus been referred to as monocyte-derived DCs (moDCs) by some authors [21] . Overall, DC ontogeny is constantly revised, with tissue-specific factors coordinating the generation of specific DC populations in a particular niche and inflammatory context.

In the context of *T. gondii* infection, it is well established that infected cDCs act as Trojan horses carrying the parasite to peripheral lymphoid organs, while infected and bystander cDCs produce IL-12 that acts to induce early IFN-γ from NK cells, and later to drive Th1 and CD8^+^ T cell responses that provide IFN-γ to activate effector molecules and mechanisms [22, 23] . In the intestinal mucosa, CD103^+^CD11b^-^ and CD103^-^CD11b^-^cDCs are the primary sources of IL-12 in response to *T. gondii* [24] .

We previously showed that Bcl3 regulates the APC function of DCs [11]. We also showed that infection with *Toxoplasma gondii* is uniformly fatal in mice globally deficient in Bcl3 (*Bcl3^−/−^*), which correlated with a defective Th1-type response [9]. We further showed that mice conditionally depleted of Bcl3 in CD11c^+^ cells, which include DCs, monocytes, macrophages and other mononuclear leukocytes, clinically phenocopied the complete knockouts and had impaired production of IFN-γ in CD4^+^ and CD8^+^ T cells, whereas innate immunity appeared to be intact [13]. In the present study, we investigated the specific role of classical dendritic cells in control of *T. gondii* infection in mice.

## Results

### Lack of Bcl3 in classical dendritic cells increases host susceptibility to fatal *T. gondii* infection

We have previously defined a critical role for the atypical IκB family member Bcl3 in host defense against fatal infection with the protozoan *Toxoplasma gondii*. Infection was uniformly fatal in that study in both complete Bcl3 KO (*Bcl3^-/-^*) mice and conditional *Bcl3^flx/flx^* mice crossed with CD11c cre mice, suggesting that resistance requires expression of Bcl3 in DCs, monocytes and/or other CD11c-expressing immune cell types. In the present study, we have investigated specific Bcl3-expressing DC subsets that may be involved by crossing *Bcl3^flx/flx^* mice with Zbtb46 cre mice, in which Bcl3 is selectively deleted in classical DCs **(Fig. S1A, B).** Wildtype C57BL/6 mice and *Bcl3^flx/flx^* mice were used as controls.

Mice were infected intraperitoneally with 15 cysts of the ME 49 strain of *T. gondii*, then were observed for 40 days for mortality and weight changes. All Bcl3 KO mice were uniformly susceptible to fatal *T. gondii* infection, whereas almost all wildtype mice survived until they were euthanized at day 40 post infection (PI) **(Fig. 1A)**. Survival curves for the *Bcl3^flx/flx^ Zbtb46 cre* and *Bcl3^-/-^* mice were superimposable. *Bcl3^flx/flx^ Zbtb46 cre^-^* littermates from the conditional knockout line showed no significant difference from wildtype mice for parasite burden over time in lung and spleen, which controls for potential effects of genetic drift and environmental differences **(Fig. S2A, B).** The body weights of all mice in all three study groups decreased after infection. However, whereas weight loss in wildtype mice stopped at approximately day 20 post infection, all mice in both knockout groups continued to lose weight until they were found dead or until preset weight loss criteria for euthanasia were met **(Fig. 1B).** These results indicate that lack of Bcl3 specifically in classical dendritic cells increases host susceptibility to fatal *T. gondii* infection.

**Figure 1:**
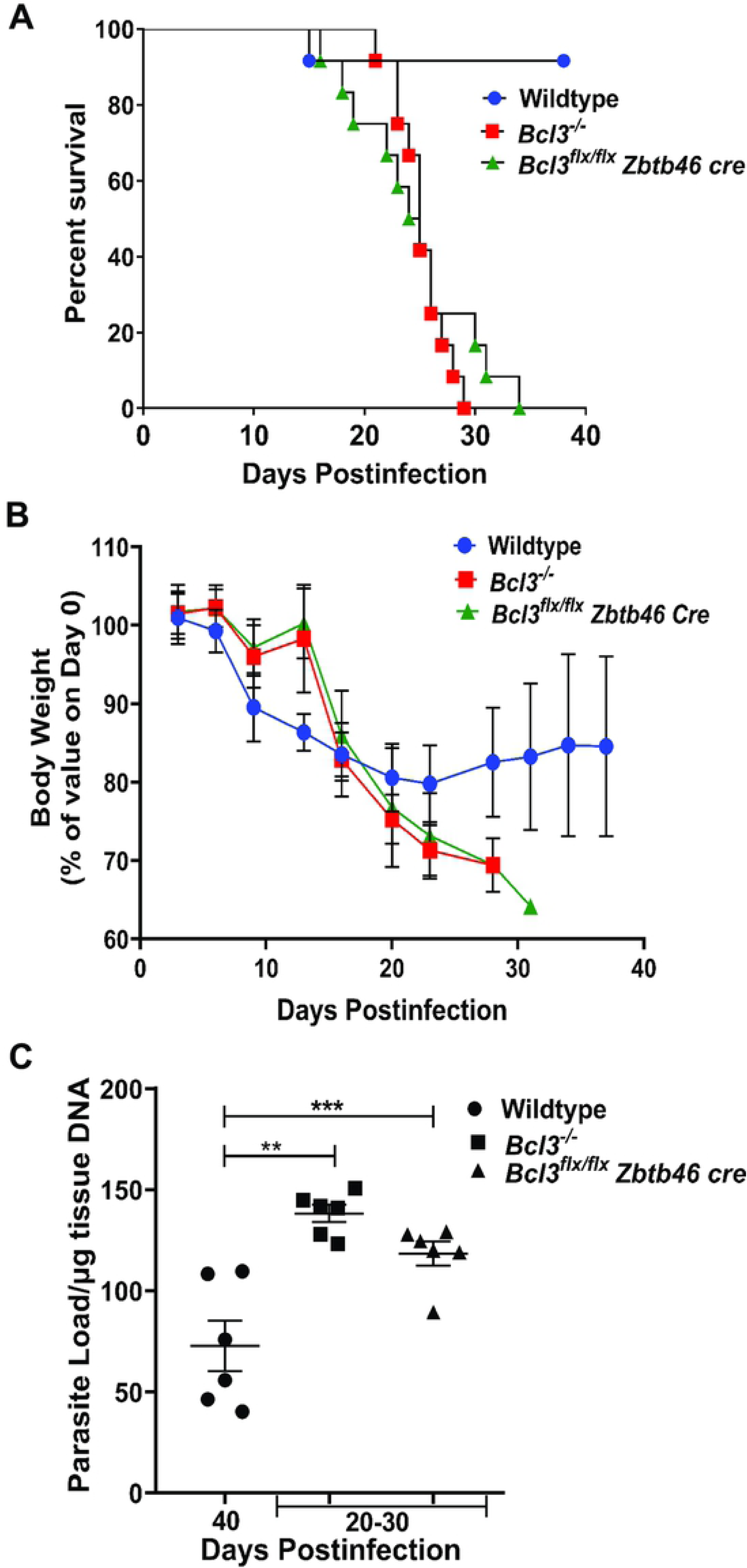
Bcl3 expression in classical dendritic cells is critical for protection against *T. gondii*. Mice with the indicated Bcl3 genotypes were infected with 15 cysts of *T. gondii* (ME49 strain) and monitored for survival (A), body weight changes (B) and brain parasite load (C). Data are summarized from 2 independent experiments with n=12 mice in each group for A and B. In C, n=6 mice in each group were selected randomly from the mice in part A. Data are shown as the mean ± SEM. The survival curve was analyzed by the log-rank Mantel-cox test. Student’s unpaired t test was used for (B) and (C). **p<0.01, ***p<0.001.

Although wild type and Bcl3 knockout mice both had increased parasite loads in all organs surveyed, the kinetic patterns varied considerably, as quantitated by measuring *T. gondii* B1 gene expression by real time PCR. Terminal brain parasite loads in both total and conditional Bcl3 KO mouse groups dying ∼20-30 days PI were significantly higher compared to parasite loads in the brains of wildtype mice sacrificed at either 20 days PI or on day 40 PI, the termination point of the experiment **(Fig. 1C** and **Fig. S1C)**. In contrast, the spleen parasite loads for both knockout groups were similar to wild type levels on day 7 PI. However, wildtype mice were then able to clear the parasite from spleen by 21 days PI, whereas for both KO groups parasite loads at day 21 PI persisted at the same levels found on day 7 PI **(Fig. 2A).** A third kinetic pattern occurred in the lung, in which parasite loads in knockout mice were similar to wildtype levels on day 7 PI, but diverged thereafter, increasing in the knockouts by day 21 PI, while remaining unaltered in wildtype mice **(Fig. S2C)**.

**Figure 2:**
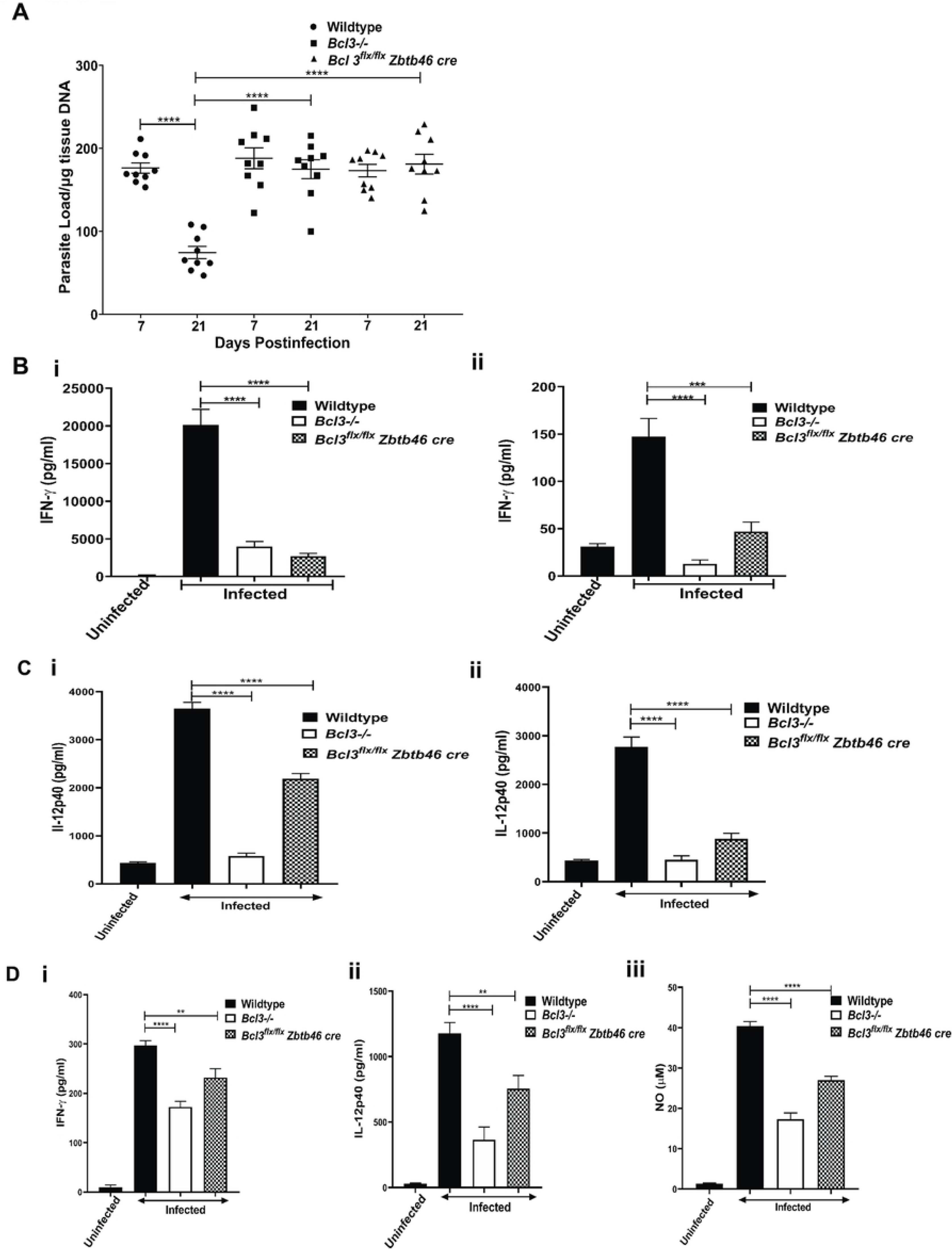
Mice lacking Bcl3 in classical dendritic cells have impaired immune responses to *T. gondii* infection and an increased parasite load. The indicated mice were infected ip with 15 cysts of *T. gondii* (ME49 strain) and assessed for spleen parasite load (A) and serum IFN-γ and IL-12 concentration (B) at 7 D PI (Bi, Ci) and 21 D PI (Bii, Cii). In panel D, splenocytes from uninfected and infected mice were stimulated ex vivo 7 days PI with STAg (5 mg/ml) for 72 hours. Supernatants were collected and IFN-γ (i), IL-12 p40 (ii) and nitric oxide (iii) levels were determined by ELISA. Data are shown as the mean ± SEM, n=3 for uninfected mice and n= 9 for all infected groups, which were pooled from 3 independent experiments. Student’s unpaired t test was used for statistical analysis. **p<0.01, ***p<0.001, ****p<0.0001.

Lung, spleen and liver from wild type and knockout mice were next evaluated histologically before and after *T. gondii* infection. In lung from infected mice, we detected perivascular and peribronchiolar 10-20 mm diameter protozoal cysts in infected lung. At 21 days PI, overall inflammation and cyst burden were both higher in all Bcl3 total KO mice and *Bcl3^flx/flx^ Zbtb46 cre* mice than in wildtype mice **(Fig. S2D)**. At baseline, spleen size was similar among *Bcl3* knockouts and wild type mice. However, 21 days after infection we observed moderate to marked enlargement of the spleen in all knockout groups compared to wildtype controls which was accompanied by hyperplasia of resident splenic lymphoid tissue but minimal inflammation. There was no evidence of significant inflammation in the liver or cyst accumulation in either spleen or liver (data not shown).

### Impaired immune responses to *T. gondii* in classical dendritic cell specific Bcl3-deficient mice

A type 1 IFN-γ-dependent immune response has previously been established as a critical factor for immunological control of *T. gondii* infection. Early innate defenses are intact even in the absence of Bcl3, as initial production of IL-12 by dendritic cells and IFN-γ by NK cells are unaffected. However, subsequent production of IFN-γ by CD4^+^ and CD8^+^ T-cells is severely compromised in complete Bcl3 knockout mice and conditional CD11c Bcl3 knockout mice. Here we confirmed this precedent and extended it by interrogating cDC Bcl3-deficient mice specifically.

First, we investigated the state of the immune system at baseline in lung and spleen in naïve uninfected wild type and knockout animals **(Fig. S3A, B)**. The content of CD11c^+^MHC II^+^ cells, which include dendritic cells, was similar for wildtype and both complete and conditional Bcl3-deficient strains in both organs. Conditional Bcl3 knockout mice had reduced frequencies of B cells and neutrophils, but only in lung, not in spleen, whereas *Bcl3^-/-^* mice had a lower frequency of B cells, neutrophils, T cells and monocytes in both organs. These results are consistent with and extend our previous report of impaired germinal center reactions associated with reduced B cell numbers in the spleen of complete *Bcl3^-/-^* mice [9].

In serum, we found that both IFN-γ and IL-12 levels were markedly increased at both 7 and 21 days after infection in both wildtype and *Bcl3^flx/flx^ Zbtb46 cre^-^* (Bcl3 sufficient) mice compared to levels in uninfected control mice **(Fig. 2B, C, Fig. S2E, F)**. In contrast, *T. gondii* infection resulted in markedly reduced serum levels of both IFN-γ and IL-12 in complete Bcl3 KOs as well as in Zbtb46 cre conditional Bcl3 KOs compared to levels induced by infection of wildtype mice.

To delineate the mechanisms underlying Bcl3-dependent Type 1 cytokine production in the model, we harvested total splenocytes from wild type and Bcl3 knockout mice 7 days PI. The total spleen content of MHC-II^+^CD11c^+^ DCs was comparable for wildtype and KO mice. After stimulation in vitro with soluble toxoplasma antigen (STAg) for 72 hr, high levels of IFN-γ, IL-12 and nitric oxide accumulated in the supernatants of cells from infected wild type mice compared to cells from uninfected wild type mice, whereas accumulation of all three mediators was significantly lower after stimulation of splenocytes from both complete and conditional Bcl3 knockout mice **(Fig. 2D).**

### Bcl3 modulates the distribution of multiple splenic dendritic cell subsets in *T. gondii*-infected mice

Next, we interrogated how DCs might be regulated at the level of gene expression by Bcl3 under inflamed conditions in *T. gondii*-infected mice. For this, infected and uninfected wildtype and *Bcl3^flx/flx^ Zbtb46 cre* mice were sacrificed 7 days PI, and CD11c^+^ splenocytes were purified and immediately processed to generate single cell RNA sequencing libraries, which were then sequenced. Using Immgen-based cell auto-annotation [25] mononuclear phagocytes (MPs) and DCs were identified and dissected into cDC1, migratory cDC1 (cDC1mig), cDC2, migratory cDC2 (cDC2mig), plasmacytoid DCs (pDCs), monocytes and splenic macrophages [26] **(Fig. S4A-C**), and this was confirmed by examining the expression pattern of the signature genes for each subset (**Fig. S4D**). To understand the transcriptomic effects of Bcl3 on splenic DCs, we selected cDC1, cDC1mig, cDC2 and cDC2mig for further analysis. Unsupervised clustering revealed 13 subpopulations among classical DCs (6 subclusters of cDC1, including cDC1mig, and 7 subclusters of cDC2, including cDC2mig) (**Fig. 3A, B**). From the differential gene expression analysis, we found common genes for cDC1 and cDC2, as well as subset-defining genes. In particular, cDC1 subpopulations expressed Cxcl9, Cst3, Irf8, Xcr1, CD24a and CD8a in common, whereas cDC2 subpopulations all expressed Ppp1r14a, Ltb, Adam23, Adgrl3, Cybb, Rgs2, Ltb, Kit, Lyz2, Zeb2, Fyb and S100 calcium-binding protein family members in common. Migratory subpopulations of both cDC1 and cDC2 expressed Ccr7, Tuba1a, Fscn1, Ccl5 and Cxcl6, which are known to be important for cell migration and lymphocyte activation. Compared to cDC1mig, cDC2mig showed much higher expression of NF-κB pathway molecules, e.g., Socs2, Traf1, Stat4, Relb and Map4k4.

**Figure 3:**
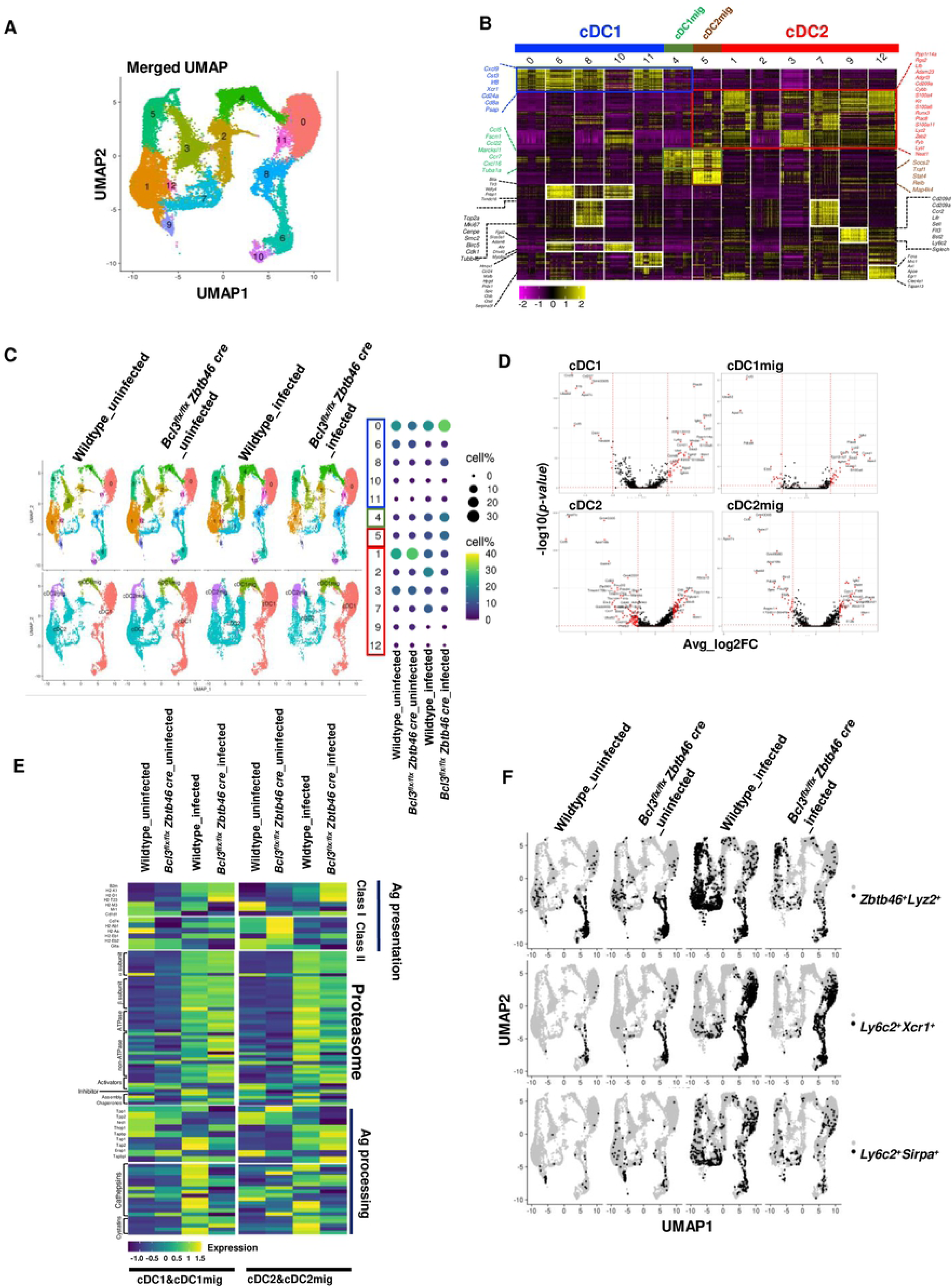
Classical dendritic cell Bcl3 deficiency distorts the distribution of dendritic cell subsets after *T. gondii* infection. Splenic CD11c+ cells were enriched by magnet-based sorting from uninfected wildtype and *Bcl3^flx/flx^ Zbtb46 cre* mice, as well as from wildtype and *Bcl3^flx/flx^ Zbtb46 cre* mice 7 day after *T. gondii* infection. The sorted cells were immediately processed for single cell RNA sequencing. (A) UMAP of splenic dendritic cell single cell data merged from all four samples (PCs = 1:30, Res = 0.3) (B) Heatmap of the top 30 cluster markers. Thirteen different clusters were re-grouped and re-ordered as cDC1 (blue bar), cDC1mig (migratory cDC1, green bar), cDC2mig (migratory cDC2, dark red bar), and cDC2 (red bar), based on shared cluster-specific genes across clusters. Colored gene name labels on either side of heatmap indicate the representative genes for cDC1 (blue box), cDC2 (red box), migratory DC (green box), and cDC2mig (dark red box). Other cluster specific genes are represented in black text. (C) UMAP split by experimental conditions, grouped by cluster (upper left panel) and grouped by DC type (lower left panel). Right panel shows the proportion (%) of each cluster in every experimental group, represented with different sized and colored circles to show how each cluster is affected by Bcl3-deficiency or *T. gondii* infection (D) Volcano plots to show genes differentially expressed between infected wildtype and infected *Bcl3^flx/flx^ Zbtb46 cre* conditions in cDC1, cDC1mig, cDC2, and cDC2mig cells significant at average log2FC > 0.5 and -log10(p-value) > 1.3. X axis denotes the average log2FC (Fold Change). Y-axis denotes the -log10 transformed p-values. (E) Heatmap showing average expression for genes associated with antigen presentation in cDC1, cDC1mig, cDC2, and cDC2mig. (F) Double-positive cells: Zbtb46 and Lyz2, Ly6c2 and Xcr1, or Ly6c2 and Sirpa were highlighted in black on split UMAP to illustrate the effects of Bcl3 deficiency and T. gondii infection. Cells with higher than 0.2 normalized gene expression value were considered as positive.

Among the cDC1 subsets, clusters 6 and 8 showed relatively higher expression of Btla and Wdfy4, which are important for peripheral regulatory T cell induction [27] and antigen cross-presentation [28]. These cells also expressed Tlr3. Cluster 8 from cDC1 and cluster 7 from cDC2 expressed cell cycle-related genes, including Top2a, Mki67, Cenpe, Birc5 and Cdk1, suggesting they may be potential DC progenitors. Clusters 9, 11 and 12 were smaller but also displayed interesting features. Cluster 9 expressed the pDC markers CD209a, Bst2, Ly6c2 and Siglech; however, the expression levels of these genes were far lower than for true pDCs, which were filtered out in the preprocessing step. Clusters 11 and 12 expressed cDC1 and cDC2 signature genes, respectively; however, both also expressed macrophage signature genes.

Next, we asked how Bcl3 deficiency during *T. gondii* infection affects the proportion of splenic DC subsets. For this, each cluster size was quantitated under the four different experimental conditions (wildtype uninfected, *Bcl3^flx/flx^ Zbtb46 cre* uninfected, wildtype infected and *Bcl3^flx/flx^ Zbtb46 cre* infected) **(Fig. 3C)**. Cluster size was not significantly different for wildtype and *Bcl3^flx/flx^ Zbtb46 cre* mice in the steady state before infection. *T. gondii* infection increased the percentage of both the cDC1mig and cDC2mig clusters to a similar extent in both wildtype and *Bcl3^flx/flx^ Zbtb46 cre* mice **(Fig. 3C, S5A)**. Migratory DCs from infected KO mice expressed more of the cytokines Il1b and Il27, and the chemotactic factors Ccr7, Cxcl9, Ccl5 and Ccl22 than migratory DCs from infected wildtype mice (**Fig. 3D**; **Table S1)**.

The proportional distribution of cDC1 sub-clusters observed in wildtype mice did not change appreciably after *T. gondii* infection, with only minor reductions in the size of clusters 6 and 10. In contrast, infection of *Bcl3^flx/flx^ Zbtb46 cre* mice resulted in increased size of subclusters 0, 8 and 11, and decreased size of subcluster 6. Infection of wildtype mice induced larger distortions of cDC2 than cDC1 subcluster distribution. In particular, the size of subclusters 1, 3, 9 and 12 decreased, whereas the size of subclusters 2 and 7 increased. Infection of *Bcl3^flx/flx^ Zbtb46 cre* mice also resulted in decreased size of cDC2 subclusters 1, 3, 9 and 12, whereas the size of cDC2 subclusters 2 and 7 did not change significantly. Moreover, larger changes in gene expression were observed in the comparison of wildtype- and Bcl3^flx/flx^ Zbtb46 cre -infected cDC2 than for the cDC1 comparison **(Fig. 3D)**.

Next, we searched the DC transcriptomic data for specific functional classes of differentially expressed genes, focusing first on genes involved in antigen presentation **(Fig. 3E, S5B, Table S2)**. cDC1 cells from both uninfected wildtype and uninfected *Bcl3^flx/flx^ Zbtb46 cre* mice expressed similar levels of genes related to antigen presentation except those related to MHC class Ib genes (H2-M3, Mr1, Cd1d), which seemed to be affected by Bcl3 deficiency even in the absence of infection. Moreover, cDC2 cells from uninfected *Bcl3^flx/flx^ Zbtb46 cre* mice had only slightly increased expression of genes involved in Class II antigen presentation (CD74, H2-Ab1, H2-Aa and H2-Eb1) from uninfected wildtype mice. In contrast, the expression level of genes involved in antigen presentation was changed dramatically in the cells from wildtype infected mice as compared to cells from uninfected mice: *T. gondii* infection increased the expression level of genes related to proteasome components, proteases (cathepsins), protease inhibitors (cystatins) and peptide delivery from the cytosol into the endoplasmic reticulum (ER) (Tap1, Tap2, Tapbp, Tapbpl) in both cDC1 and cDC2 cells. Interestingly, the level of genes encoding peptidases (Tpp1, Tpp2, Nrd1, Erap1), which are known to be required for the generation of most MHC class I-binding peptides, were decreased after infection. Class I pathway related genes (B2m, H2-K1, H2-D1) were increased or slightly increased in wild-type infected cDC1 and cDC2 cells, respectively, whereas Class II pathway related genes were overall downregulated in cDC2 cells from infected wildtype mice as compared to cells from uninfected mice. Striking changes from the comparison between wildtype and Bcl3 KO cells from infected mice were observed in the category of antigen processing genes: cDC1 cells showed reduced levels of genes associated with peptide delivery. Additionally, most cathepsins and cystatins were significantly decreased in both cDC1 and cDC2 cells. Together, these changes suggest a reduced capacity for cross-presentation in Bcl3 KO cDC1 cells compared to wildtype even though the expression level of MHC class I genes was intact or slightly increased in KO cells. The overall expression level of genes involved in proteasome assembly was unaltered by Bcl3 knockout in the context of *T.gondii* infection.

Since DC Bcl3 deficiency has previously been reported to accelerate apoptosis of bone marrow– derived DCs during Ag presentation to T cells, and to impair DC survival in the context of inflammatory conditions [11], we next interrogated genes associated with apoptosis and inflammation in the data set, visualizing the results by tSNE transformation **(Fig. S5C, D**, **Table S2)**. As expected, *T. gondii* infection per se induced dramatic changes in expression of apoptosis and inflammation-related genes. Cluster 1, a subcluster of cDC2, showed the most significant differences in distribution of these functional classes of differentially expressed genes under both uninfected and infected conditions.

Next, we examined the expression patterns of genes encoding transcription factors with particular attention to Zbtb46 because it is a hallmark of classical DCs and because we used its promoter to delete Bcl3 in cDCs using cre/lox technology. We identified unique DC subpopulations that coexpressed both Zbtb46 and LysM, a transcription factor previously thought to be exclusively expressed in monocytes and macrophages. These dual-positive or hybrid cells were mainly found in cluster 6 in both uninfected wildtype and *Bcl3^flx/flx^ Zbtb46 cre* mice; however, after infection of wildtype mice they became widely distributed among all the DC clusters. Compared to infected wildtype mice, the frequency of Zbtb46^+^LysM^+^ dual positive cells were markedly reduced in infected *Bcl3^flx/flx^ Zbtb46 cre* mice **(Fig. 3F, upper panel, and Fig. S5E, F)**. A second monocyte/macrophage gene Ly6C and the classical DC gene XCR1 were also coexpressed in some clusters but these hybrid cells were only marginally detected in uninfected mice. Infected wildtype mice showed a significant increase in this population, which in contrast was diminished in frequency in infected *Bcl3^flx/flx^ Zbtb46 cre* mice **(Fig. 3F, middle panel, and Fig. S5E, F)**. Finally, we found that cells coexpressing Ly6C and the cDC2 marker gene Sirpa were significantly more frequent in infected wildtype mice compared to infected conditional KO mice **(Fig. 3F, lower panel, and Fig. S5E, F**). Thus, the data suggest that *T. gondii* infection may induce differentiation of unique DC subpopulations with dual monocyte/macrophage and DC characteristics at the transcriptomic level.

To understand the physiological significance of the hybrid cells, we decided to perform functional enrichment analysis (Ingenuity Pathway Analysis). For this, Zbtb46 and Lyz2 dual positive cells were sorted out from the four different experimental conditions **(Fig. S6 A)**. The expression level of Lyz2 in these dual positive cells was much lower than in splenic macrophages, which reaffirms these cells are not monocytes/macrophages **(Fig. S6 B)**. We also found that *T.gondii* infection induced Lysozyme M expression exclusively in wildtype mice, however, classical DC lacking Bcl3 do not show an infection induced upregulation in its expression **(Fig. S6 C)**. DEGs from comparisons between hybrid and nonhybrid cells from uninfected or infected conditions, or comparisons between wildtype and Bcl3 KO hybrid cells from uninfected or infected conditions were used as an input. Unlike hybrid cells in steady state, which seem to be quiescent **(Fig. S6 D)**, hybrid cells in *T.gondii* infection clearly showed stronger immune cell response related pathways, like Th1 pathway, TCR signaling, CD28 signaling, iNOS signaling, unfolded protein response, phagosome formation and etcs, suggesting their potential contribution to anti-intracellular bacterial immune responses **(Fig. S6 E)**.

Comparison between uninfected vs infected hybrid cells showed increased signaling through canonical IFN signaling pathway, activation of IRF, dendritic cell maturation, NF-kB signaling and oxidative phosphorylation in infected hybrid cells **(Fig. S6 F)**. Further analysis between wildtype and Bcl3 KO cells from infected mice revealed unregulated up-regulated phagosome formation and unfolded protein response, indicative of their superior antigen processing and presentation capacity **(Fig. S6 H)**. Importantly, cell cycle checkpoint related pathways were increased in KO cells. There were no significant changes between uninfected hybrid cells from wildtype and Bcl3 KO counterparts by IPA **(Fig. S6 G)**. Altogether, Zbtb46 and Lyz2 dual positive cells are immunologically more activated type of DCs emerged in response to the pro-inflammatory cytokines e.g. interferons in *T.gondii* infected condition where Bcl3 plays a key role to maintain the hybrid cells and their anti-intracellular bacterial capacity.

### Immunophenotypic characterization of two novel DC subsets, icDC1 and icDC2, regulated by Bcl3 expression and *T. gondii* infection

Based on the single cell RNA seq data, we revisited the identity of DC subsets in the spleen and lung 7 days PI using cell surface markers and flow cytometry. We identified two subsets of cDC1 and cDC2, which we have designated icDC1 and icDC2 due to their high frequency in lung and spleen from *T. gondii*-infected mice compared to uninfected mice. Both subsets were found in wild type mice as well as in complete and conditional Bcl3 knockout mice. In addition to the conventional DC markers CD11c, MHC II, CD24, CD8 and CD11b, icDC1 are defined by co-expression of XCR1 and Ly6C, which is a prototypic marker for the monocytic/macrophage lineage and inflammatory DCs, and icDC2 are defined by co-expression of CD11b and Ly6C. Both icDC1 and icDC2 also express CD64, another macrophage marker **(Gating strategy Fig. 4A, and Fig. S7A)**. The difference in frequency of these subsets in infected versus uninfected mice defined by flow cytometry aligned with the difference in frequency as defined by the RNA seq data.

**Figure 4:**
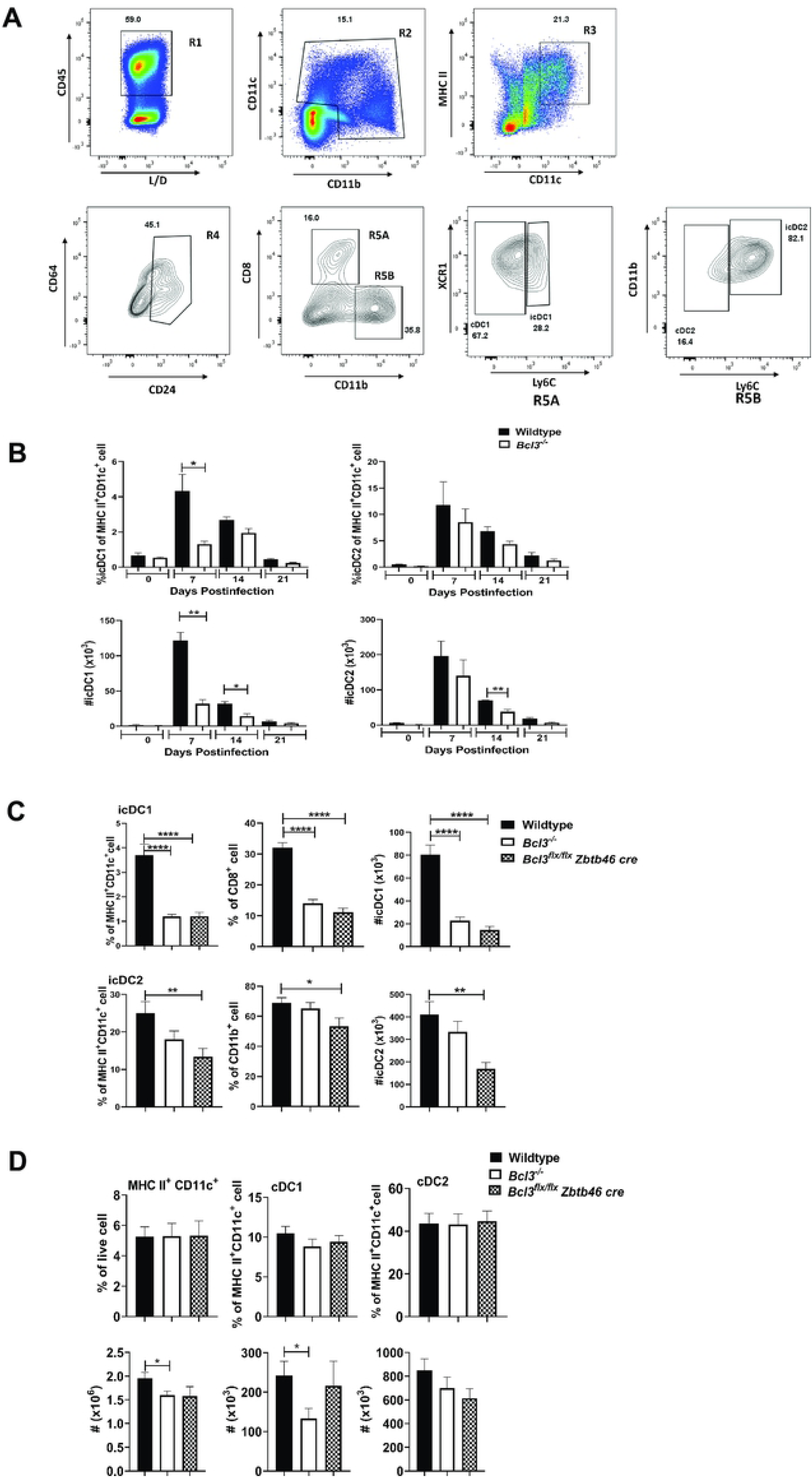
Generation of hybrid conventional DCs in *T. gondii*-infected spleen. Mice were infected with 15 cysts of *T. gondii* (ME49 strain) and DC phenotyping of splenocytes was performed 7 days post infection. Novel infection-associated DC subsets were designated icDC. (A) Gating strategy for dendritic cell subsets. (B) Time course for accumulation of the indicated DC subpopulations in wildtype and *Bcl3^-/-^* mice. (C and D) Bcl3-dependent distribution of splenic icDC (icDC1 and icDC2) (C) and cDC (cDC1 and cDC2) (D) subsets. The Bcl3 genotype code is shown in the upper right of each panel. Data are summarized as the mean ± SEM of n = 6 mice/group pooled from 2 independent experiments. Student’s unpaired t test was used for statistical analysis. *p<0.05, **p<0.01, ****p<0.0001.

As early as 7 days after infection, dramatic but transient increases in both the frequency and number of icDC1 occurred in both spleen and lung in wild type mice, however this increase was markedly reduced in infected total Bcl3 KO mice **(Fig. 4B, and Fig. S7B)**. icDC2 levels were also increased by infection in spleen and lung; however, the peak levels were similar for both wild type and total Bcl3 KO mice. Moreover, the increase in spleen was transient whereas in lung the increase was more sustained, persisting as late as 21 days post infection. In infected *Bcl3^flx/flx^ Zbtb46 cre* mice, induction of both icDC1 and icDC2 cells was reduced on day 7 PI in both lung and spleen compared to results in wildtype control mice **(Fig. 4C and Fig. S7C**). In contrast, total cDC1 and cDC2 content in spleen and lung on day 7 post *T. gondii* infection was not affected by specific deletion of Bcl3 in cDCs using the Zbtb46 cre or in total KO (**Fig. 4D and Fig. S7D**).

These findings align with the RNA seq data where XCR1^+^Ly6C^+^ and Ly6C^+^Sirpα^+^ co-expressing cells are regulated by Bcl3 expression in Zbtb46^+^ classical DCs.

### Defective antigen presentation and T cell priming in mice selectively deficient in Bcl3 in classical dendritic cells

To delineate the functional role of Bcl3 specifically in classical dendritic cells in the model, we compared antigen presentation and cytokine production by immune cells from wildtype and *Bcl3^flx/flx^ Zbtb46 cre* mice. To assess antigen presentation, MACS-purified CD11b^+^ and XCR1^+^ cells from naïve animals were pulsed with ova peptide or were infected in vitro with ova-expressing *T. gondii* tachyzoites, then were cocultured with OT-I CD8^+^ T cells, which were assessed for proliferation **(Fig. S8A)**.

CD11b^+^ cells from wildtype and *Bcl3^flx/flx^ Zbtb46 cre* mice stimulated similar levels of OT-I T cell proliferation. In contrast, XCR1^+^ DCs from wild type mice had far superior antigen-presenting activity (both for ova peptide and for naturally processed ova protein from ova-expressing *T. gondii*) compared to XCR1^+^ DCs from *Bcl3^flx/flx^ Zbtb46 cre* mice, whose activity was close to background for the assay **(Fig. 5A, B)**. This proves that XCR1^+^ DCs are pivotal for cross-presentation of ova antigen and that Bcl3 regulates this function in this subset of DCs.

**Figure 5:**
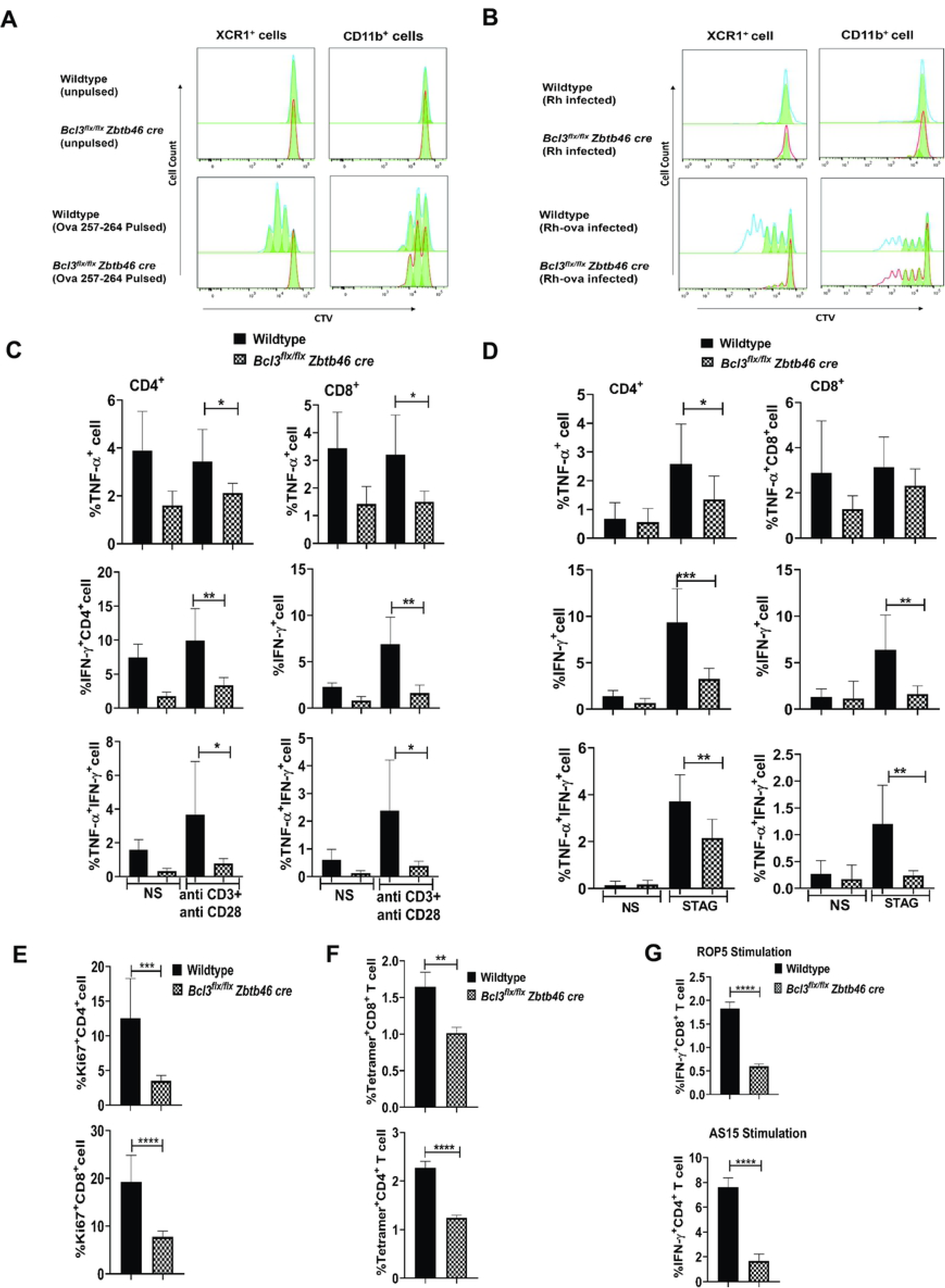
Bcl3 is pivotal for antigen presentation by XCR1^+^ conventional dendritic cells and antigen-specific T cell responses against *T. gondii*. (A, B) Defective antigen-specific T cell proliferation. Splenic XCR1^+^ and CD11b^+^ cells were isolated from naive wildtype and *Bcl3^flx/flx^ Zbtb46 cre* mice. Cells were pulsed with class I (Kb)- restricted OVA peptide 257-264 for 3 hours (A) or infected with ova-expressing *T. gondii* (Rhcontrol and Rh-ova) for 24 hours (B). Finally, they were cocultured with Cell tracer violet-stained OT-I T cells for an additional 72 hours. T cells were analyzed by flow cytometry and the proliferation profile for CD8^+^ T cells was determined. Data are representative of 2 independent experiments. (C-G) Defective antigen-specific T cell function. (C, D, E) Wildtype and *Bcl3^flx/flx^ Zbtb46 cre* mice were infected with 15 cysts of *T. gondii* (ME49 strain), then splenocytes were isolated 18 days later and stimulated with plate-bound anti-CD3 and soluble anti-CD28 for 6 h (C) or with STAg for 72 h (D). Intracellular IFN-γ and TNF-α in CD4^+^ (left) and CD8^+^ (right) cells were measured by flow cytometry. (E) Ki67 staining was assessed for unstimulated CD4^+^ and CD8^+^ T cells from infected spleen. Representative plots are summarized as the mean ± SEM of n = 8 mice/group pooled from 3 experiments. Student’s unpaired t test was used for statistical analysis. (F, G) Splenocytes were harvested from wildtype and *Bcl3^flx/flx^ Zbtb46 cre* mice 3 weeks post infection. *T. gondii*-specific CD4^+^ T and CD8^+^ T cell responses were measured by MHC class I/II tetramer staining (F) and by intracellular IFN-γ staining after in vitro restimulation with T. gondii-specific peptide for 4 hours (G). Representative plots are summarized as the mean ± SEM; n= 10 mice/group pooled from 3 experiments. Student’s unpaired t test was used for statistical analysis. *p<0.05, **p<0.01, ***p<0.001, ****p<0.0001.

To assess cytokine induction by DCs in the model, we isolated splenocytes at day 18 PI from wildtype and *Bcl3^flx/flx^ Zbtb46 cre* mice, a timepoint when the adaptive immune response has begun in response to the infection. The T cells were stimulated ex vivo with plate-bound anti-CD3 and soluble anti-CD28. Intracellular IFN-γ and TNF-α levels were significantly reduced in both CD4^+^ and CD8^+^ T cells from infected *Bcl3^flx/flx^ Zbtb46 cre* mice compared to wild type mice. We also observed a significantly lower frequency of multifunctional IFN-γ^+^TNF-α^+^ CD4^+^ and CD8^+^ T cells among activated splenocytes from infected mice deficient in Bcl3 in classical DCs as compared with activated T cells from infected wildtype mice **(Fig. S8B, C; Fig. 5C)**.

Similarly, when splenocytes were stimulated ex vivo with STAg to induce a *T. gondii*-specific response, we found that IFN-γ levels were dramatically reduced in CD4^+^ and CD8^+^ T cells from infected *Bcl3^flx/flx^ Zbtb46 cre* mice compared to T cells from infected wild type mice. STAg-stimulated TNF-α levels were also reduced in splenic T cells from infected *Bcl3^flx/flx^ Zbtb46 cre* mice compared to wildtype controls, but only in CD4^+^ T cells, not in CD8^+^ T cells, whereas dual IFN-γ^+^TNF-α^+^ cells were reduced in both CD4^+^ and CD8^+^ T cell compartments after STAg stimulation of splenic T cells from infected *Bcl3^flx/flx^ Zbtb46 cre* mice compared to wildtype controls **(Fig. S8B, C; Fig. 5D)**. Consistent with these cDC Bcl3-dependent cytokine responses, in response to STAg stimulation of splenocytes, both CD4^+^ and CD8^+^ T cells from infected wildtype mice had significantly increased evidence of proliferation, as determined by Ki67 staining, compared to cells from infected *Bcl3^flx/flx^ Zbtb46 cre* mice **(Fig. S8D, Fig. 5E)**.

To further examine the role of cDC Bcl3 in the *T. gondii*-specific immune response, we infected wildtype and *Bcl3^flx/flx^ Zbtb46 cre* mice, isolated splenocytes 3 weeks PI and subsequently stained them for *T. gondii* tetramer-positive T cells. Both Tetramer^+^CD4^+^ and Tetramer^+^CD8^+^ T cell frequencies were much higher in splenocytes harvested from infected wildtype mice than in splenocytes from infected *Bcl3^flx/flx^ Zbtb46 cre* mice, providing evidence for a pronounced antigen-specific Bcl3-dependent response **(Fig. S8E, Fig. 5F).** Splenocytes from day 21 PI were also stimulated with the *T. gondii* MHC-I-restricted peptide AS15 and the MHC-II-restricted peptide ROP5 for 4 hours. Intracellular IFN-γ generation was significantly higher in peptide-activated CD4^+^ and CD8^+^ T cells from wildtype mice compared to T cells from *Bcl3^flx/flx^ Zbtb46 cre* mice **(Fig. S8F, Fig. 5G)**.

### Bcl3 promotes the development of classical dendritic cells

After establishing a role for Bcl3 in classical DCs in the *T. gondii*-specific adaptive immune response, we next investigated whether Bcl3 might regulate cDC development. For this, we exploited an established bone marrow differentiation protocol in which a combination of Flt3L and NOTCH2 signaling is used for terminal differentiation of classical DC1, which are specialized for cross presentation. In this approach, murine bone marrow hematopoietic progenitors are cocultured with DL1 (NOTCH2 ligand)-expressing fibroblasts (OP9DL1 cells) in the presence of Flt3L. Unlike OP9 cells (DL1 negative fibroblasts), when cDC1 are differentiated in the presence of NOTCH signaling, they generate bona fide cDC1 with proper phenotypic markers (CD8α^+^, Dec 205^+^) and better T cell cross priming potential. We used this approach to differentiate bone marrow cells in vitro from uninfected wildtype and *Bcl3^-/-^* mice **(Fig. S9A, Scheme)**. We found that coculture with OP9 fibroblasts generated a small proportion of cells expressing CD24 and XCR1, which are markers for cross-presenting cDC1s, however this is not affected by Bcl3 deficiency or IFN-γ addition to the culture system. **(Fig. S9B)**. However, when differentiated in the presence of DL1 (OP9DL1 fibroblasts), wildtype bone marrow developed cells expressing both cDC1 markers. In contrast, bone marrow from Bcl3-deficient mice cocultured with OP9-DL1 fibroblasts generated cells with lower levels of expression of CD24 and XCR1 **(Fig. 6A).**

**Figure 6:**
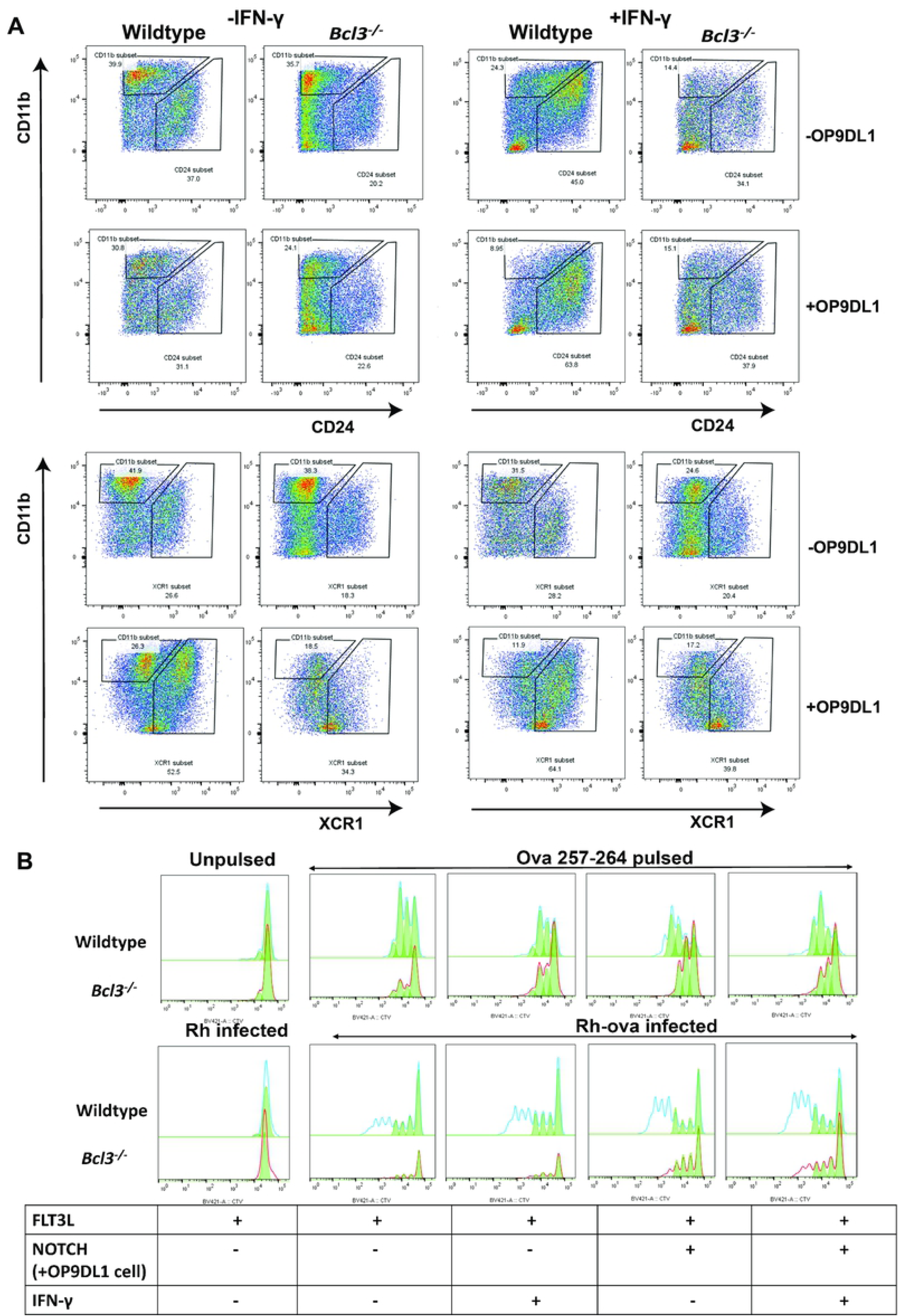
Bcl3 deficient bone marrow-derived DCs fail to differentiate into potent antigen presenting cells. (A) Skewed DC differentiation in Bcl3-deficient cells. Bone marrow cells were isolated from wildtype and *Bcl3^-/-^* mice and differentiated as described in Supplementary Fig. 8A with and without IFN-γ and OP9DL1 cells as defined above and to the right of each plot, respectively. Expression of the indicated DC surface markers was examined after 7-9 days post differentiation. Data are representative of 3 independent experiments. (B) Defective antigen presentation by Bcl3-deficient DCs. Differentiated BMDC from wildtype and *Bcl3^-/-^* mice were used as APC and were either pulsed with class I (Kb)-restricted OVA peptide 257-264 for 3 hours or infected with ova-expressing *T. gondii* for 24 hours. Finally, the cells were cocultured with Cell tracer violet-loaded OT-I T cells for an additional 72 hours. T cells were analyzed by flow cytometry and CD8^+^ T cell proliferation was determined. Data are representative of 2 independent experiments.

IFN-γ is also a critical regulator of DC differentiation and is strongly induced during *T. gondii* infection. Therefore, we tested its ability to modulate cDC1 development in our coculture system as a function of Bcl3 expression. For this, we added exogenous IFN-γ to a 7 Day old coculture for an additional 2 days. IFN-γ alone in the absence of NOTCH2 signaling but in the presence of Flt3L signaling did not affect the level of cDC1 differentiation observed in cocultures of bone marrow from wildtype or *Bcl3^-/-^* mice. However, in the presence of NOTCH2 signaling, IFN-γ significantly increased CD24- and XCR1-expressing cells differentiated from wildtype bone marrow, but not from Bcl3-deficient bone marrow **(Fig. 6A, right panel)**. Thus, IFN-γ and NOTCH2 cannot compensate for Bcl3 deficiency for the generation of immunophenotypically defined cDC1s. To test Bcl3 regulation of cDC1 function in this system, we either pulsed the bone marrow-differentiated DCs with ova peptides or infected them in vitro with ova-expressing *T. gondii*. Subsequently, they were cocultured with CD8^+^ OT-I T cells whose proliferation was monitored. In the absence of NOTCH2 signaling, ova stimulation only slightly increased APC function, with wildtype DC activity greater than activity for *Bcl3^-/-^* -derived DC **(Fig. 6B, upper panel)**. In the presence of NOTCH signaling, cross presentation is substantially improved in the system for wild type DCs, but not for Bcl3-deficient DCs **(Fig. 6B, lower panel)**.

## Discussion

In the present study, we have demonstrated that mice lacking the NF-κB regulator Bcl3 in cells expressing Zbtb46, a selective immune cell marker of cDCs, uniformly succumb 3-5 weeks after intraperitoneal infection with *T. gondii*. Infected *Bcl3^flx/flx^ Zbtb46 cre* mice failed to clear the parasite in brain, spleen and lung and had impaired Th1 immune responses, with reduced production of the critical macrophage-activating cytokine IFN-γ from antigen-specific CD4^+^ and CD8^+^ T cells in the adaptive phase of the infection, as early as 18 days PI. These results extend our previous report of *T. gondii* outcome in global Bcl3 knockout mice and mice conditionally deficient in Bcl3 in cells expressing CD11c, which include all subsets of DCs as well as multiple other immune cells, including neutrophils, NK cells, NKT cells, B cells, monocytes and macrophages [9, 13]. At the clinical level, we found that survival kinetics were the same for *Bcl3^flx/flx^ Zbtb46 cre* mice as for total Bcl3 KO mice, defining Bcl3 expression specifically in cDCs as a key survival factor. While our study does not address or exclude a role for Bcl3 in *Toxoplasma* susceptibility in other Bcl3-expressing CD11c^+^ cell types, it did allow us to focus our attention on the specific mechanistic role of cDC Bcl3 in the model.

Dendritic cells act as a key player in host defense against *Toxoplasma* infection. After initial encounter by the parasite, *Toxoplasma*-infected and bystander DCs produce IL-12 along with macrophages and monocytes, with neutrophils contributing to a lesser extent. IL-12 is pivotal for generating a Th1 immune response with early innate production of IFN-γ by natural killer (NK) cells in the acute phase and later by CD4^+^ T cells in the adaptive phase and CD8^+^ T cells in the chronic phase [17, 29, 30] . Inflammatory mediators such as nitric oxide are under direct control of IFN-γ [31]. Bcl3 deficiency impairs IFN-γ-inducible nitric oxide generation, thereby allowing prolonged parasite survival in the peripheral organs and brain.

*T. gondii* enters the dendritic cell by active invasion and is sequestered in specialized compartments called parasitophorous vacuoles [32]. Several studies have demonstrated that the parasite can be detected by the immune system using the MHC I-mediated endogenous pathway of professional APCs in cross priming CD8^+^ T cells in the chronic phase of the infection [33] XCR1^+^ classical DCs are mediators of cross presentation in mouse and human [34]. In this regard, our previously published work on Bcl3 function in DCs was limited to analysis of 7-9 day in vitro GM-CSF-stimulated bone marrow-derived CD11b^+^ CD11c^+^ dendritic cells (BMDCs), in which Bcl3 deficiency reduced BMDC maturation and survival after ovalbumin antigen/adjuvant challenge as well as BMDC priming of OT-II CD4^+^ T cells and cross priming of OT-I CD8^+^ T cells both in vitro and, in adoptive transfer experiments, in vivo [11]. Also, our previously published immunologic analysis of the effects of Bcl3 deficiency in the toxoplasmosis challenge model was limited to demonstrating that IFN-γ production from NK cells during the early innate immune response was normal in *T. gondii*-infected global *Bcl3^-/-^* and *Bcl3^flx/flx^ CD11c cre mice* (CD11c Cre-driven deletion) and that accumulation of IFN-γ-producing CD4^+^ and CD8^+^ T cells in spleen was reduced at day 18 PI [6].

Our present results extend these precedents in two ways. First, we directly examined the Bcl3 dependence of *T. gondii* antigen-specific T cell responses in primary splenocytes. Here we employed *T. gondii*-specific tetramer staining and intracellular IFN-γ generation following *T. gondii*-specific peptide stimulation ex vivo. Proliferation of these activated T cells was monitored by Ki67 staining. And second, we used single cell RNAseq technology to describe heterogeneous cDC subsets at the molecular level that are induced by *T. gondii* in the spleens of infected mice in a cDC Bcl3-dependent manner.

With regard to antigen presentation and T cell activation, we found that 1) *T. gondii* infection resulted in much lower serum levels of both IFN-γ and IL-12 in cDC Bcl3-deficient mice; 2) the production of IFN-γ, IL-12 and nitric oxide induced by STAg stimulation in vitro of whole splenocytes harvested at day 7 PI was markedly reduced in cDC Bcl3-deficient mice; 3) Bcl3 was required for normal cross presentation of antigen by XCR1^+^ cDC1s to OT-I CD8^+^ T cells (both for exogenous ova peptide or naturally processed ova protein from ova-expressing *T. gondii* after infection); however presentation by CD11b^+^ cells, consisting of cDC2, monocytes, monocyte-derived DC and macrophages remained essentially intact; 4) at day 18 PI, a timepoint when the *T. gondii*-specific adaptive immune response is well-established, splenic CD4^+^ and CD8^+^ T cells from cDC Bcl3-deficient mice had reduced levels of intracellular IFN-γ and TNF-α and a lower frequency of multifunctional IFN-γ^+^TNF-α^+^ cells after stimulation *ex vivo* with anti-CD3 and anti-CD28, as well as impaired STAg-induced CD4^+^ and CD8^+^ T cell proliferation and IFN-γ production; and 5) in splenocytes harvested at day 21 PI from cDC Bcl3-deficient mice there were reduced frequencies of *T. gondii* tetramer-positive CD4^+^ and CD8^+^ T cells and reduced intracellular IFN-γ-positive CD4^+^ and CD8 ^+^ T cells after stimulation with the *T. gondii* MHC-II-restricted peptide AS15 and the MHC-I-restricted peptide ROP5. Together, the data provide evidence that Bcl3 expression in cDCs is critical for antigen-specific T cell responses, including responses to *T. gondii* antigens in vivo. Moreover, the data confirm that XCR1^+^ cDC1s are pivotal for cross-presentation of antigen and demonstrate that Bcl3 regulates this function in the context of *T. gondii* infection. How Bcl3 exclusively regulates antigen presentation in cDC1s and not antigen presentation in CD11b^+^ cells, consisting of monocytes, macrophages and cDC2s (including MoDCs) is an open question revealed by our study worthy of future investigation.

Dendritic cells are extremely heterogeneous in terms of their phenotype and functions. They form organ-specific subsets with diversity in surface markers, migratory patterns, localization, and cytokine production. The development of diverse populations of DCs is differentially regulated by differentially expressed transcription factors and cytokines [18]. Furthermore, several studies have been reported describing the emergence of nonconventional dendritic cells with distinct immunological roles. Inflammatory cDC2 acquire CD64, a macrophage marker and express cDC1-specific Irf8 in lungs of mice infected with the single-stranded RNA virus pneumonia virus of mice (PVM), a virus closely related to human respiratory syncytial virus (RSV) [35]. Type I IFN-induced CD64^+^ cDCs have also been described in the context of *Listeria* infection [36]. Even under steady state conditions, classically defined cDC2s contain a discrete population of apparent monocyte-derived cells capable of DC function, including cross-presentation [37]. Further, tumor immunotherapy using a PTEN inhibitor (vanadate drug VO-OHpic) has been reported to induce the generation of Batf3-dependent, CD103^+^ Ly6C^+^ cross-presenting cells arising from an immature monocytic precursor present in the peripheral MDSC pool [38]. In our study, we revisited the concept of dendritic cell heterogeneity under a similar IFN-γ induced inflammatory milieu in *T. gondii* infection. We could demonstrate from scRNA seq analysis that *T. gondii*-infected spleen contains a hybrid DC with dual expression of Zbtb46 and LysM, the signature transcription factors for classical DCs and monocytic cells, respectively. To our knowledge, this is the first report for the existence of such a unique subpopulation. We found a small population of these cells in naïve uninfected wildtype mice which cluster closely with cells phenotypically defined as cDC1. However, after infection, these cells were distributed in all subclusters of cDC1s and cDC2s. Since the lifespan of DCs is 10-14 days and since they are constantly replenished by BM precursors, we speculate that special precursor cells are generated and expanded under inflammatory pressure. Zbtb46 is expressed by the immediate precursor of cDCs (precDC1 and precDC2), but not in early and intermediate DC progenitors [39]. Hence, the Bcl3-specific defect cannot be traced back to MDP or CDP differentiation.

We confirmed the existence of these ‘hybrid’ cells by flow cytometry as CD11c^hi^MHC II^hi^CD24^+^CD8^+^XCR1^+^CD64^+^Ly6C^+^ cells and CD11c^hi^MHC II^hi^CD24^+^CD11b^+^/Sirpa^+^CD64^+^Ly6C^+^ cells, which we refer to as inflammatory cDC1 (icDC1) and inflammatory cDC2 (icDC2) cells, respectively. Interestingly, both icDC1 and icDC2 are reduced in both frequency and absolute number in both lung and spleen from infected *Bcl3*^-/-^ mice and *Bcl3^flx/flx^ Zbtb46 cre* mice. How Bcl3 is regulating the generation of these subsets is beyond the scope of the present work; however, we speculate that development of icDC1 and icDC2 during infection might provide a Tip-DC-like function in otherwise conventional DCs that might support T cell priming and TNF-α and nitric oxide generation [40].

Previous studies on host transcriptomics have revealed a general host-pathogen interaction in an in vitro *T. gondii* infection model [41] and in cat intestine [42]. To our knowledge, our study provides the first single cell transcriptomic investigation of dendritic cell Bcl3 in an experimental murine model of Toxoplasmosis. Since antigen presentation is a professional function of classical DCs and is regulated by Bcl3, we focused on genes known to affect this function. Overall, MHC expression was not affected by Bcl3 deficiency, whereas proteosomal genes and genes involved in antigen processing were most highly dependent on Bcl3 for upregulation during infection. We also identified Bcl3-dependent genes involved in apoptosis, cell migration and inflammation, which showed distinct expression patterns in mice with different genotypes and under different experimental conditions.

cDC development from BM progenitors is mainly driven by Flt3L. Interestingly, NOTCH signaling along with Flt3L results in exclusive generation of cDC1 with distinct phenotypic markers and enhanced T cell priming capacity [43]. We found that in vitro differentiation of BM cells from Bcl3-deficient mice to XCR1^+^CD24^+^ cDC1 in response to NOTCH/Flt3L stimulation was defective and could not be rescued by addition of exogenous IFN-γ. Thus, Bcl3 deficiency may cause an intrinsic defect in pre-cDC1. Additionally, these in vitro generated cDC1 are less potent in presenting antigen (both ova peptide and ova protein in the context of *T. gondii*) to CD8^+^ OT-I T cells than wildtype control cDC1.

In conclusion, our study establishes a role for Bcl3 in development of cDCs in the context of *T. gondii* infection and inflammation, including confirmation of novel inflammatory cDC subsets defined by transcriptomic and immunophenotypic criteria. We have extended our previous studies of Bcl3 in toxoplasmosis by assigning a specific role of cDC antigen cross presentation and CD8^+^ T cell activation. Finally, our single cell RNAseq data suggest that the effect of Bcl3 deficiency in cDC gene expression in the model includes major effects on antigen processing genes, which provide new and testable hypotheses for future studies of the functional role of Bcl3 in toxoplasmosis.

## Materials and Methods

### Ethics statement

All animal handling procedures and experiments were approved by the NIAID Animal Care and Use Committee (protocol LMI-23E) and were conducted in accordance with all relevant institutional guidelines.

### Mice

*Bcl3^-/-^* mice [9] and *Bcl3^flx/flx^* mice [11] were generated in our laboratory and previously described. Zbtb46 cre mice was a kind gift from Dr. Michel Nussenweig [39]. *Bcl3^flx/flx^ Zbtb46 cre* mice (Bcl-3 knockout in classical dendritic cells) were generated by crosses of *Bcl3^flx/flx^* and *Bcl3^KO/flx^* mice carrying the Zbtb46-driven Cre recombinase transgene. All the Bcl3 sufficient controls were littermates. *Bcl3^-/-^* (Taconic line 74), WT (Taconic line) and OT-I mice (Taconic line 175) were purchased from Taconic Biosciences (Germantown, NY, USA). All mice were based on the C57BL/6 background. All mice were housed in NIAID Institute facilities.

### Parasite

The ME49 strain of *T. gondii* was maintained in wildtype C57BL/6 mice by intraperitoneal injection (15 cysts/mice). After 30 days mice were sacrificed, and brain cysts were isolated and reinfected into naïve animals. Rh and Rh-ova-Td tomato tachyzoites were maintained in Hs27 cells (human foreskin fibroblasts). Confluent monolayer cells were infected with the tachyzoite form of the parasite at an M.O.I of 1:10. After 72-96 hours, the cells burst due to the parasite load. The tachyzoites were collected and reinfected into fresh cells.

### Cells

Hs27 cells were maintained in DMEM medium supplemented with 10% FCS. OP9 and OP9-DL1 (expressing NOTCH ligand DL1) cells (macrophage derived ESC from bone marrow) were maintained in Alpha minimum essential medium supplemented with 2.2 g/L sodium bicarbonate and 20% FCS.

### Infection and survival kinetics

For experimental infections, mice were inoculated i.p. with an average of 15 cysts/animal and monitored for survival and weight change.

### Genomic DNA isolation and B1 gene PCR

Organ sections were collected from infected mice at the indicated times and genomic DNA was isolated using the DNeasy Blood and tissue kit (Qiagen, Cat. No: 69504). The B1 gene from *T. gondii* was amplified and organ parasite load was determined using a standard curve [44]. 500 ng of genomic DNA was used in a SYBR green- (Cat no. A25776, Applied Biosystem, Thermo Fisher Scientific Waltham, MA USA) based real time PCR reaction in Quantstudio 3 (Applied Biosystems) using the Standard curve with Melt protocol.

**Forward primer:** F 5’-CTC CTT CGT CCG TCG TAA TAT C-3’

**Reverse primer:** R 5’-TGG TGT ACT GCG AAA ATG AAT C-3’

Cycling conditions: UDG activation, 50°C, 2’; Initial denaturation, 95°C, 10’ followed by 40 cycles of 95°C, 30’’; 62°C, 40’’; and 72°C, 1’; Final extension, 72°C, 5’; Melt curve, 95°C (1.6°C/sec), 15’’; 60°C (1.6°C/sec), 15’’.

### Bcl3 PCR in classical DC

Spleen cells were isolated from *Bcl3^flx/flx^* and *Bcl3^flx/flx^ Zbtb46 cre* mice and CD11c cells were sorted using CD11c MicroBeads (Militenyi Biotec, Cat# 130-125-835) according to the manufacturer’s instructions. Next Zbtb46^+^ cells (cDC) and Zbtb46-cells (non cDC, CD11c^+^Zbtb46^-^) were FACS sorted after intracellular staining. T cells (Militenyi Biotec, Cat# 130-095-130) were isolated as controls. PCR was performed using the primer sets for floxed and KO alleles:

KO Forward: 5’ GCGCCGCCCCGACTGAC 3’

Floxed Forward: 5’ CGTCCCCAGAGCCCGCAACCAC 3’

Reverse (common): 5’GGGCCTCTCAACCTCTTTCCTA 3’

Zbtb46 cre PCR was performed according to the Jackson Laboratory protocol (Stock Number: 028538)

### Serum Cytokines

Mice were sacrificed at day 0, day 7 and day 21 post infection and blood was collected. Serum was isolated by centrifugation and stored at -80°C. Serum was diluted if necessary and IFN-γ (1:100) and IL-12 (1:50) were measured by ELISA using a BD Biosciences kit, (BD OptEIA™ Mouse IFN-γ (AN-18) ELISA Set, Cat#551866; BD OptEIA™ Mouse IL-12 p40 ELISA Set, Cat # 555165) according to the manufacturer’s protocol.

### Ex vivo stimulation

Total splenocytes were isolated from uninfected mice and mice 7 days pi. 4 x 10^6^ cells were stimulated ex vivo by 5 μg/ml of soluble toxoplasma antigen (STAg) for 72 h. Culture supernatants were collected and extracellular cytokines were measured using an ELISA kit from BD Biosciences. Nitric oxide was measured using the Griess reagent system from Invitrogen (Cat #G7921).

### Histology

Organs were immersion-fixed in 10% buffered formalin and embedded in paraffin blocks. Sections were stained with hematoxylin and eosin (H&E) and examined by light microscopy.

### Ag presentation

For measuring direct antigen presentation, 5 x 10^4^ Splenic DC (XCR1^+^ DC, isolated using the Anti-XCR1 MicroBead Kit (Spleen), mouse, Cat no. 130-115-721, Miltenyi Biotech or CD11b^+^ cells isolated using CD11b MicroBeads UltraPure, mouse, Cat no. 130-126-725, Miltenyi Biotech) or BMDCs were stimulated for 3 h using Ova peptide 257–264 (SIINFEKL) (Cat no. AS-60193-5, AnaSpec, Fremont, CA, USA). For measuring antigen cross presentation, the cells were infected in vitro with Ova-expressing Rh tachyzoites (Ova-Rh-td-tomato) at an MOI of 1:10 for 24 h. The cells were washed and cocultured with 2.5 x 10^5^ Cell Tracer Violet (Cat no. C34557 A, Invitrogen)-loaded CD8^+^ OT-I cells (isolated using CD8a^+^ T Cell Isolation Kit, mouse, Cat no. 130-104-075, Miltenyi Biotech) for 72 h. Subsequently the proliferation profile of the OT-I cells was determined by Flow cytometry.

### Tetramer staining and intracellular cytokine determination

Splenocytes were isolated from mice 18 days PI and stained with ROP5-MHC-I tetramer and AS15-MHC-II tetramer (*Toxoplasma gondii* specific tetramers, synthesized by the NIAID, NIH tetramer core facility, Atlanta, GA, USA) [45, 46] at room temperature and 4°C, respectively, for 1 hour. Dead cells were stained with Live/Dead Aqua (Cat no. L34966, Life Technologies Corporation, Eugene, OR, USA) along with surface antibodies. To determine intracellular cytokine production from CD4^+^/CD8^+^ T cells, splenocytes were isolated from 18-day infected mice and stimulated with plate-bound anti-CD3ε (Clone 145-2C11, 2 μg/mL) and soluble αCD28 (Clone 37.51, 1 μg/mL) (both from BioXcell, West Lebanon, NH, USA) for 6 h, or STAg (5 μg/ml) for 72 h, or the Toxoplasma specific peptides AS15 and ROP5 (custom made, Genscript) for 4 hours. Cells were cultured in the presence of a protein transport inhibitor cocktail (Cat no. 00-4980-93, eBioscience; Thermo Fisher Scientific, Carlsbad, CA, USA) for the last 4 hr. Cells were stained with Live/Dead Aqua and cell surface markers, fixed and permeabilized and finally stained for intracellular cytokines at 4°C using antibodies listed in Table 1 (Supplementary information).

### In vitro BMDC differentiation

Single cell suspensions were generated from wildtype and Bcl3 KO BM cells. The cells were suspended in DMEM medium supplemented with 10% FCS, 1% L-glutamine, 1% sodium pyruvate, 1% MEM-NEAA and 1% penicillin/streptomycin, 55 mM 2-mercaptoethanol and Flt3L (100 ng/ml) (Cat no. RP-8665, Invitrogen, Thermo Fischer Scientific) and cultured at 37°C in a humidified atmosphere at 5% CO2. On day 3, the cells were transferred to a single well containing a monolayer of mitomycin C (Cat no. 50-07-7, Millipore Sigma, Merck, Darmstadt, Germany)-treated OP9/ OP9-DL1 cells or were kept unaltered. At Day 7, the cells were supplemented with murine rIFN-γ (40 μg/ml) (Recombinant Murine IFN-γ, Catalog Number:315-05, Peprotech) or were kept unaltered. At Day 9, all the cells were harvested and used for DC phenotyping by Flow cytometry or were used further in the antigen presentation assay described above.

### CD11c^+^ splenocyte isolation for single-cell RNA sequencing and library preparation

Single cell suspensions of splenocytes were enriched for CD11c^+^ cells using CD11c MicroBeads (Militenyi Biotec, Cat# 130-125-835) according to the manufacturer’s instructions. The downstream procedures of single-cell RNA-seq library preparation from the CD11c-enriched single cells and library sequencing were performed by the Single Cell Analysis Facility (SCAF) of the National Cancer Institute (NCI) Center for Cancer Research. scRNA-seq libraries were prepared using the Chromium Single Cell 3’ Reagent Kits v3.1 (10X Genomics; Pleasanton, CA, USA) according to the manufacturer’s instructions. Generated libraries were sequenced on an Illumina NextSeq 2000 instrument, followed by de-multiplexing and mapping to the mouse genome (mm10: refdata-gex-mm10-2020-A) using cellranger (10X Genomics, version 4.0.0).

**This dataset is available at GEO Series accession number GSE193532 (https://www.ncbi.nlm.nih.gov/geo/query/acc.cgi?acc=GSE193532).**

### scRNA-seq data analysis

Gene expression matrices were generated using cellranger (10X Genomics, version 4.0.0) and the raw matrices were further processed using the Seurat package (4.0.1) [47] in R (version 4.0.5). For quality control, the following categories were excluded from the analysis: (i) genes expressed by fewer than 3 cells; (ii) cells with lower than 200 or more than 6000 genes detected; (iii) cells in which >20% of unique molecular identifiers (UMIs) were derived from the mitochondrial genome. To align shared cell populations across datasets, multiple experimental single-cell datasets were integrated using the “anchoring” strategy to remove batch effects. This involved combining multiple datasets and normalizing them and finding highly variable features individually using “NormalizeData” and “FindVariableFeatures” functions respectively from Seurat package. Common features that repeatedly vary across datasets (determined using “FindIntegrationAnchors” function) were used as integration anchors for integrating multiple datasets using “IntegrateData” function from Seurat. Integrated dataset was scaled and clustered using Louvain algorithm (resolution = 0.3). Dimensionality reduction was performed using Principal Component Analysis (PCA, n=30), t-stochastic neighboring embedding (t-SNE) and Uniform Manifold Approximation and Projection (UMAP) for visualization. Transcriptomic mouse datasets from the Immgen database [26] were used for reference-based cell type annotation using SingleR (v1.0.5) [25]. From the auto-annotated data, only cells identified as dendritic cells or macrophages were selected and data was re-normalized and re-clustered for finer analyses. Cluster size was depicted as its proportion within a group and the significance of the difference between the proportion of cells in clusters between groups was calculated using scProportionTest (v1.0.0) package. Differentially expressed genes (DEGs) between clusters were calculated using “FindAllMarkers” or “FindMarker” functions by Wilcoxon rank sum test (default) from Seurat. To maximize the visualization of DEGs between clusters or experimental groups, we used the “AverageExpression” function within Seurat. Gene clustering for heatmap visualization was performed by hierarchical clustering (the “hclust” function from stats package (v3.6.2) using either ‘complete’ or ‘ward.D2’ methods). To overcome the dropout effect in single cell data, we used the MAGIC package (2.0.3) [48] with the default setting (knn = 5, decay = 1) in supplementary figures. To sort out dual-positive or hybrid cells by the expression of *Zbtb46*, *Lyz2*, *Ly6c2*, *Xcr1* and *Sirpa*, the normalized gene count matrix was extracted from the Seurat object using the “GetAssayData” function and the cells with > 0.2 normalized gene expression value considered as positive. Functional enrichment analysis was performed through Ingenuity Pathway Analysis.

### Immune cell staining

Cells from naïve or infected mice at the indicated time intervals were collected from lung or spleen and separate panels of antibodies were used for immunophenotyping as listed in Table1 (Supplementary information).

### Statistical analysis

Data were recorded as the mean ±SEM. Differences between groups were analyzed by unpaired, two-tailed Student’s t-tests. Results with a p value of 0.05 or less were considered significant (Prism; GraphPad Software). Survival studies were analyzed by the log-rank Mantel-Cox test. The number of independent data points (n) and the number of independent experiments is stated in figure legends.

## Acknowledgements

This research was supported by the Intramural Research Program of the National Institute of Allergy and Infectious Diseases, National Institutes of Health, Bethesda, MD. We thank Dragana Lj Jankovic for kindly providing the *Toxoplasma gondii* (ME49) cysts, Rh Tachyzoites and Hs27 cells; Ian Moore for helping with the Histological studies; Michel Nussenzweig for kindly providing the Zbtb46 cre mice; Christopher A. Hunter for providing the ova-Rh-td-tomato tachyzoites; and Michael Kelly for help with scRNA library preparation and sequencing. Support from the CCR Single Cell Analysis Facility was funded by FNLCR Contract HHSN261200800001E. This work utilized the computational resources of the NIH HPC Biowulf cluster (http://hpc.nih.gov). Finally, we thank the NIH tetramer facility for preparing the Toxoplasma tetramers.

## Supporting Information caption

**S1 Fig:** Genotyping and brain parasite load

**S2 Fig:** Immune response and histopathology

**S3 Fig:** Immune cell distribution in naïve mice

**S4 Fig:** Preprocessing of scRNAseq data and identification of cells predicted by unsupervised clustering

**S5 Fig:** DC distribution is significantly influenced by Bcl3 deficiency in the context of *T. gondii* infection

**S6 Fig:** Characterization of Hybrid cell from scRNA seq results in spleen

**S7 Fig:** Generation of hybrid conventional DCs in T. gondii-infected lung

**S8 Fig:** Schematics of Antigen presentation assay and Dot plots of Intracellular cytokine staining

**S9 Fig:** Schematics of invitro BMDC differentiation and immunophenotyping

